# Enlarged Ingroup Effect: How a Shared Culture Shapes In-Group Perception

**DOI:** 10.1101/2020.06.12.148601

**Authors:** Paola Rigo, Bindiya L. Ragunath, Marc H. Bornstein, Gianluca Esposito

## Abstract

Increasing levels of migration and constant redefinition of a ‘sense of belonging’ characterize modern societies. Thus, social perception of people from different ethnicities as in-group or out-group members is influenced by a shared culture that might go beyond ethnicity. Using functional magnetic resonance imaging, we aim to study how sharing a common culture changes the social perceptions of in-groups and out-groups. We presented same- and different-race faces to young adults living in an integrated multicultural society. Same- and different-race faces were primed by images of environmental context that promotes identification with the participants’ ethnicity or a common shared culture.

We found that same and different-race faces recruit similar brain networks only when associated with an environmental context, which promoted identification with a common shared culture. These results support a possible emergent phenomenon in multicultural societies that we call enlarged multi-ethnic in-group effect, which may form the basis of a potential new way to categorize oneself and others in terms of membership.

## Introduction

Due to rapid urbanization and globalization resulting in increasing levels of migration (Kærgård, 2010), many countries today have residents of various ethnic backgrounds (Dijkstra, Geuijen, & De Ruijter, 2001; Holtug, 2016; Mahmud & Jahan, 2013; Keong, 2013). A challenge for such multicultural societies is to understand complex social interactions to ensure social cohesion and harmony (Dijkstra et al., 2001; Rosenthal & Levy, 2010). People make sense of the complex social world through social categorization (Derks, Stedehouder, & Ito, 2014; Kaul et al., 2012; Van Bavel & Cunningham, 2010). People perceive themselves and other people they encounter as members of social categories based on salient social categories, such as, ethnicity, gender, age, and nationality, (Derks et al., 2014; Van Bavel & Cunningham, 2010; Kurzban, Tooby, & Cosmides, 2001), some of which are automatically encoded (Kaul et al., 2012), and which shape the early processing of visual features of others (Derks et al., 2014).

In multicultural societies, the negative aspects of ethnic diversity are addressed at communal and political levels (Gijsberts, Van Der Meer, & Dagevos, 2011; Keong, 2013). Rosenthal and Levy (2010) summarized a multicultural approach that proposes social categorization, such as race and ethnicity, should be given attention and understood in order to develop knowledge and respect for other groups. According to these authors, multiculturalism is characterized by three central aspects: acknowledgment of differences between ethnic groups; appreciation of each group’s contributions to society; and emphasis on groups maintaining their own culture and traditions (Rosenthal & Levy, 2010). Enhanced social interaction and integration can lead to the development of a common shared culture across multi-ethnic groups that all members can identify with, in addition to their ethnic identity.

On one hand, there is a tendency to divide the social world into ‘us’ and ‘them’ (Shkurko, 2012; Van Bavel & Cunningham, 2010), and on the other, in-group and out-group categorization can be fluid, context-dependent, and mediated by its relevance for self-definition in terms of personal or social identity (Turnet et al., 1994). In fact, people can be members of various groups simultaneously or categorize others as in-group and out-group members depending on the more relevant categorization in the ongoing social situation. For example, some categorization criteria include the individual’s goals, expectations, desires, and beliefs important for self-identification (Hehman et al, 2011; Katsumi & Dolcos, 2018; Wheeler and Fiske, 2005; Turner et al., 1994, Cheon & Esposito, 2020). Social perceptions of people as in-group or out-group members are influenced by ethnicity, formal and informal social roles that drive relations between minority and majority ethnic groups in daily life, as well as many other psychological factors (i.e., facial features, emotional reactions, and cognitive aspects) (Elfenbein & Ambady, 2002; Soto & Levenson, 2009).

Faces can communicate information quickly and efficiently (Hugenberg & Wilson, 2013; Caria et al., 2012; Esposito et al., 2014; Esposito et al. 2015; Venturoso et al. 2019). Research suggests that neural responses to faces present an internal reflection of in-group and out-group perceptions. Much research on face perception and recognition were aimed at understanding in-group and out-group perceptions (e.g., Bavel, Parker, & Cunningham, 2008), mainly because faces provide important social cues critical to social categorization (e.g., sex, race, age), emotional evaluation, and intentions. Investigating neural responses to in-group and out-group social perceptions has gained ample attention (for review see Adolphs, 2001; Kubota, Banaji, & Phelps, 2012; Shkurko, 2012). Neural substrates involved in out-group social cognition using faces include visual attention networks such as the fusiform gyrus (FG); amygdala in self-regulatory processes; and social areas such as anterior cingulate cortex (ACC), temporoparietal junction (TPJ), and the superior temporal sulcus (STS).

Research suggests intergroup conflict and bias can be reduced by shifting categorizations such that former out-group members are included in the in-group (Gaertner & Dovidio, 2000). Members who belong to double in-groups (e.g., same race/same university) are preferred over members of partial in-groups (e.g., same race/different university), which in turn are preferred over members of double out-groups (e.g., different race/different university) (Crisp & Hewstone, 1999; Hehman et al., 2011). Furthermore, EEG findings showed that in face-recognition of double in-group members, compared to partial in-group or double-outgroup, participants’ brain responses were sensitive to contextual information beyond ethnicity (e.g., Hehman et al., 2011). Double in-group faces evoke a larger event-related potential (ERP) component, followed by faces of partial in-groups, indicating more significant neural processing in the attention networks (Hehman et al., 2011).

In this study, we propose that face processing of same-race and other-race ethnicities will differ based on the cultural context shared by people. Recruited participants were citizens living in Singapore, a multicultural social environment where citizens identify with their ethnic as well as their national identity, and whose multicultural stance accords with the three central aspects in multiculturalism postulated by Rosenthal & Levy (2010) (Salleh, 2017; Straits Times, 2017).

In this study, we investigated the effect of ethnic and shared cultural contexts on same-race and other-race face perception in young adults of Chinese ethnicity. Participants underwent fMRI scanning while viewing same-race and other-race faces preceded by an ethnic context (e.g., Chinese temple before a Chinese face or Taj Mahal before an Indian face), or a shared cultural context pictures (e.g., famous Singapore national monument/building).

Since our participants live with people of specific ethnicities with whom they share everyday life (typical ethnicities), we considered two types of out-groups in the present study: atypical and typical other-race faces. We postulated that neural responses to other- (out-group) and same-race faces (in-group) faces would differ in environmental contexts that promote identification with one’s ethnicity consistent with literature. In contrast, identifying with a shared culture (i.e., cultural context) would result in similar neural responses to other- and same-race faces, potentially indicating an automatic and spontaneous sense of membership to a multi-ethnic group. When primed by cultural context, we argue that *typical other-race faces,* but not *atypical other-race faces* (shared culture or enlarged in-group effect), and *same-ethnicity faces* (same-race in-group effect) will recruit similar brain regions.

We first hypothesize that typical other-race faces (Indian), when preceded by ethnic context (race effect), increase the brain activation in regions typically associated with out-group face categorizations (fusiform gyrus and visual cortex), self-regulatory and approaching/avoidant behaviors (amygdala, IFG). As a control condition for out-group faces, we included non-typical ethnic faces (Arabic, Caucasian).

In contrast, we then postulated that both typical other-race faces and same-race faces, when preceded by shared cultural context as opposed to ethic context, would increase brain activations in regions associated with personal identity, self, empathic aspects, and pro-social behaviors. Specifically, the midline brain structures (PCC-Cuneus MPFC), superior-inferior parietal cortex, lingual gyrus, STG, and inferior frontal cortex (Xu et al., 2009; Molenberghs 2013; Scheepers et al., 2013; Katsumi & Dolcos, 2017). Findings will be discussed with relevance to existing literature along with implications.

## Methods

### Participants

A total of 48 participants (50% females) aged between 21 to 26 years (*M* = 22.81, *S.D*. = 1.17) took part in this study. All participants were recruited from Nanyang Technological University (NTU), Singapore. All participants were Singaporean Chinese who had not travelled out of Singapore for more than 2 months over the past 6 months from the time of their experimental session. They were also screened to ensure right-handedness, normal or corrected-to-normal vision, and no history of psychological or neurological disorders. No female participants were pregnant. Participants had to refrain from alcohol, nicotine, and caffeine consumption 24 hours prior to the scan session. This study was approved by the NTU IRB (Protocol 2017-01-029). Written informed consent was obtained from all participants. All data are available at this URL: https://doi.org/10.21979/N9/IC672L

### Experimental Task

To measure effects of context (common shared cultural context (*CulCon*) or ethnic context (*EthCon*), on brain responses to in-group faces (*IF*) and out-group faces (*OF*)), we asked participants to view images of faces primed with *CulCon* or *EthCon* during the fMRI scan. Since the participants were all Singaporean Chinese, ethnically, *IF* consisted of Chinese faces. *OF* consisted of a common typical ethnicity present in Singapore, Indian faces, and non-typical ethnicities of Singapore, Arab and Caucasian faces. Faces of four ethnicities were categorized into typical ethnicities (Chinese and Indian faces) of Singapore and non-typical ethnicities (Arabic and Caucasian faces). Typical ethnicities were further categorized into *IF* (Chinese faces) and *OF* (Indian faces). *CulCon* comprised famous buildings/monuments of Singapore (e.g., Merlion, Esplanade) and *EthCon* contained famous buildings relevant to each ethnicity (e.g., Taj Mahal for Indian ethnicity, Chinese temple for Chinese ethnicity, Eifel Tower for Caucasian ethnicity, and the Sheik Zayed Mosque for Arabic ethnicity). All images were presented in grayscale. All faces were female and were masked with a grey round window so that only facial features were visible (no hair or neck were visible). Each trial started with a fixation cross displayed between 7 – 10 seconds, followed by a contextual picture (context condition) displayed for 2 seconds, then followed by an image of a face for 4 seconds (Figure 1). The task consisted of a total of 32 trials. The faces and their preceding contextual picture stimuli were randomized for each participant. The duration of the fixation cross was also randomized. To assess the effect of *EthCon*, only congruent *EthCon* to ethnic faces were considered (e.g., Indian faces only preceded by Taj Mahal, Caucasian faces only preceded by Eiffel Tower).

**Figure 1.**
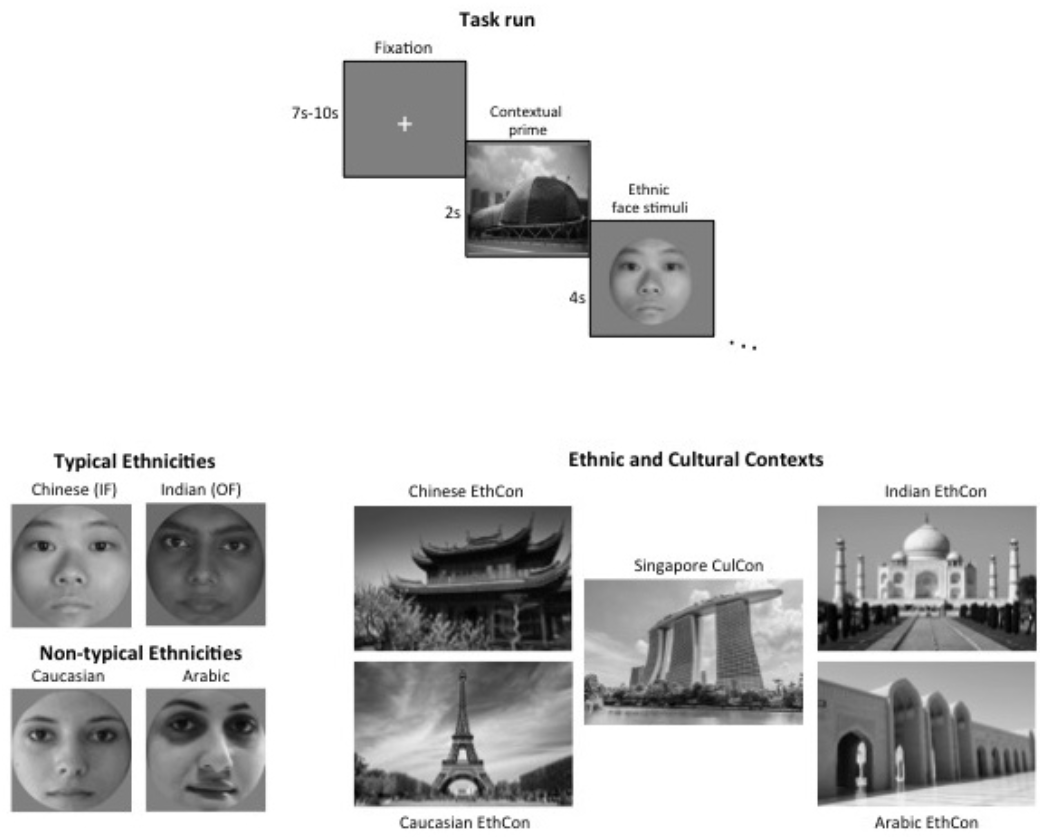
Task paradigm.

### Procedure

Prior to their testing session, eligible participants were briefed about the study and underwent a MRI safety briefing before they were scanned. Participants were also reminded of the voluntary nature of the study. The experimental task took place during the fMRI scan, where the images were displayed on a screen, and participants viewed them through a mirror positioned at their eye level on the head coil. Participants were instructed to simply view the images displayed to them during the scan. Anatomical scans were also obtained. After the scan session, participants were debriefed about the specific aims of the study and were provided monetary reimbursement of S$50.

### MRI and fMRI Data acquisition

Whole brain neuroimages were collected using a Siemens Magnetom Prisma 3-Tesla MRI Scanner with 64-channel head coil. A high-resolution T1-weighted MPRAGE sequence (192 slices; TR 2300ms; TI 900ms; flip angle 8 degrees; voxel size 1mm) was obtained to serve as an anatomical reference. Subsequently, functional images were obtained using a gradient Echo Planar Imaging (EPI) sequence with 36 axial slices (271 slices; slick thickness 3mm with no inter-slice gap), and the following parameters: TR 2000ms; TE 30ms; flip angle 90 degrees; FOV 192 × 192 mm; voxel size 3mm; interleaved. External head restraint (i.e. neck padding) was used to minimize head movement during the scan.

### fMRI Data analysis

fMRI analysis was conducted using Statistical parametric mapping (SPM12; (http://www.fil.ion.ucl.ac.uk/spm/software/spm12) on the Matlab 2017b platform. Functional image preprocessing consisted of the following steps: Discarding the first two volumes of the functional time series, correcting for head movement, and then co-registering the T1 anatomical image to the mean of the realigned functional images carried out functional image preprocessing. Functional images were then normalized to the Montreal Neurological Institute (MNI) stereotaxic standard space and finally smoothed spatially (9-mm full-width half-maximum Gaussian kernel) and temporally (cut-off period 256 seconds). Analytic design matrices were constructed for each participant, which modeled the onsets and duration of the face image from each trial as epochs convolved with a hemodynamic response function.

General linear modeling (GLM) was performed to assess the effects of ethnic context (*EthCon)* and culture context (*CulCon)* amongst typical ethnicities in Singapore (Chinese and Indian faces) and non-typical ethnicities in Singapore (Arabic and Caucasian faces). A total of four conditions were modeled as separate regressors at individual level (1^st^ Level) analysis: typical ethnic faces in *CulCon*, typical ethnic faces in *EthCon*, non-typical ethnic faces in *CulCon*, and non-typical ethnic faces in *EthCon*. Specific weight vectors were denoted to derive contrast images for second-level (group) mixed effects analysis using a general linear model (GLM). The contrasts of interest were:

1.1 *typical faces in CulCon vs. EthCon* to investigate the effect of culture and ethnic contexts on typical ethnicities; and
1.2 *non-typical faces in CulCon vs. EthCon* to assess the effect of cultural and ethnic contexts on non-typical ethnic faces.

In the 2nd level random-effect analysis, one sample t-tests were conducted to assess group effects on the contrast images from the 1st level. The t-tests indicated whether observed differences in the abovementioned contrasts (1.1 and 1.2) differed significantly from zero (Holmes and Friston, 1998) and allowed for inferences across participants.

A second GLM was performed to further assess the effect of cultural and ethnic context amongst *IF* and *OF* of typical ethnicities. A total of four conditions were modeled as separate regressors; *IF* in *EthCon*, *OF* in *EthCon*, *IF* in *CulCon*, and *OF* in *CulCon*. Specific weight vectors were denoted to derive contrast images for group level (2nd Level), mixed-effect analysis. The contrasts of interest were

2.1 *IF in CulCon vs. EthCon* to compare neural responses of in-group faces primed with cultural context to in-group faces primed with ethnic context;
2.2 *OF in CulCon vs. EthCon* to compare neural responses of out-group faces primed with cultural context to out-group faces primed with the ethic context;
2.3 *OF vs. IF in CulCon* to assess how neural responses to *OF* and *IF* differed in the cultural context; and lastly,
2.4 *OF vs. IF in EthCon* to assess how neural responses to *OF* and *IF* differed when primed with ethnic context.

Additionally, two other SPM(t) contrasts were created to assess how different the contrasts *OF in CulCon vs. IF in CulCon (2.3)* were from the contrast of *OF in EthCon vs. IF in EthCon (2.4).*

Finally, through the conjunction analysis, we assess the similarity between contrasts (2.3) and (2.4).

For all analyses, a threshold significance of p< 0.05 was set for resulting SPM(t) with a cluster-wise (k = 10) family-wise error rate (FWE) correction for multiple comparisons (Logan & Rowe, 2004). Whole brain analyses were performed.

## Results

### Neural Responses to Typical (1.1) and Non-typical (1.2) faces in Cultural compared to Ethnic Context

Comparing faces of typical ethnicities present in Singapore (Chinese and Indian) in the cultural (CulCon) *versus* the ethnic (EthCon) contexts, we found significant activated clusters as follows:

– *Typical faces in CulCon > typical faces in EthCon:* Higher activations in the right superior temporal sulcus (STS), left middle temporal gyrus (MTG), right supramarginal gyrus (SMG), right visual association area (VAA), left VAA extending to the lingual gyrus (LG), and the right primary sensory area (PSA) comprising of the postcentral gyrus (postCG).
– *Typical faces in EthCon > typical faces in CulCon:* Higher activation of the fusiform gyrus (FG) and middle occipital gyrus (MOG).

Comparing faces of non-typical ethnicities (Arab and Caucasian) in the cultural CulCon *versus* the EthCon contexts, we found significant activated clusters as follows:

– *Non-typical faces in EthCon > non-typical faces in CulCon:* the only activation seen was in the right fusiform gyrus (Table 1, Figure 2).

**Table 1.**
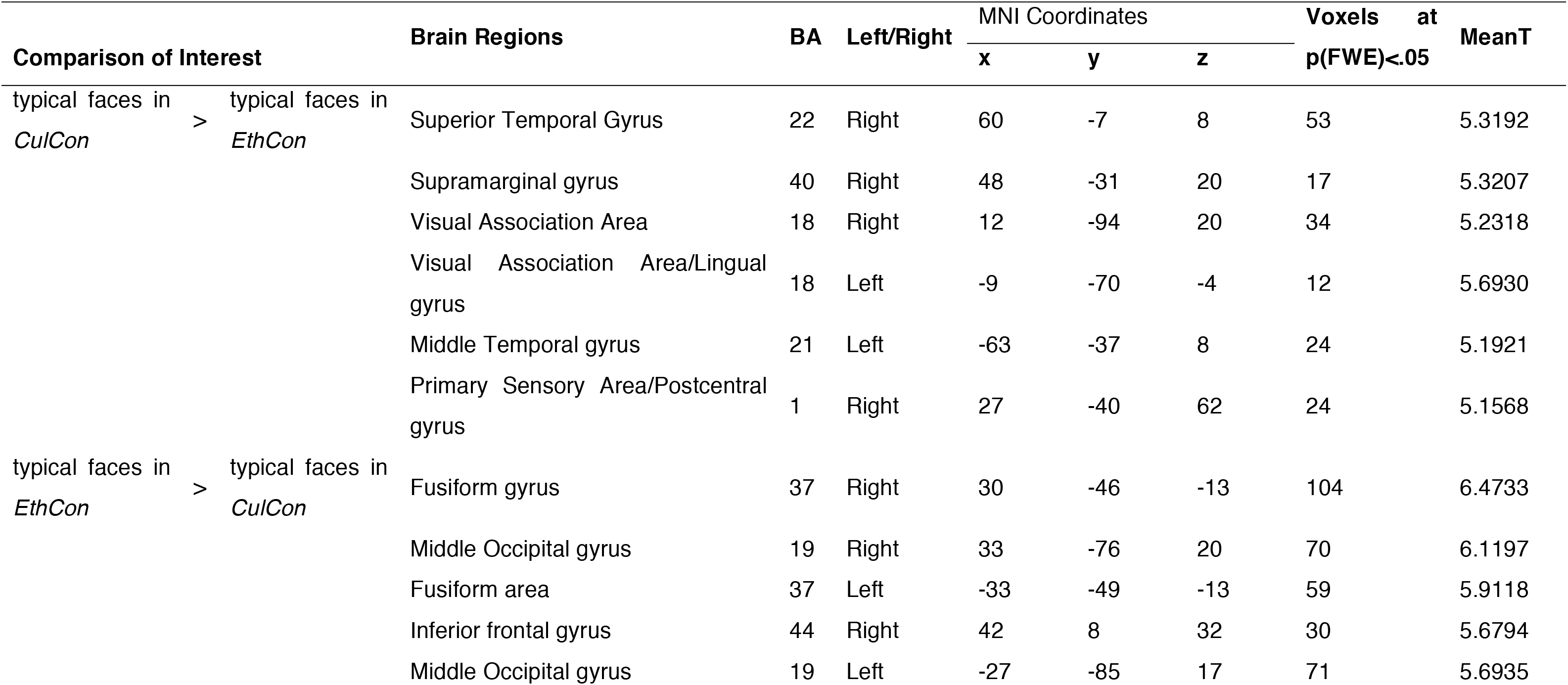
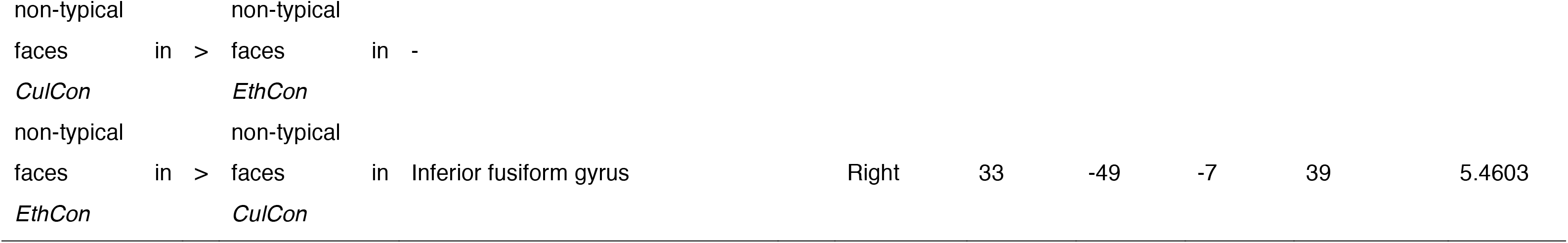
Sites of significant neural activation (p < 0.05, FWE correction) related to the contrasts typical faces in *CulCon* vs. typical faces in *EthCon* and non-typical faces in *CulCon* vs. non-typical faces in *EthCon*.

**Figure 2.**
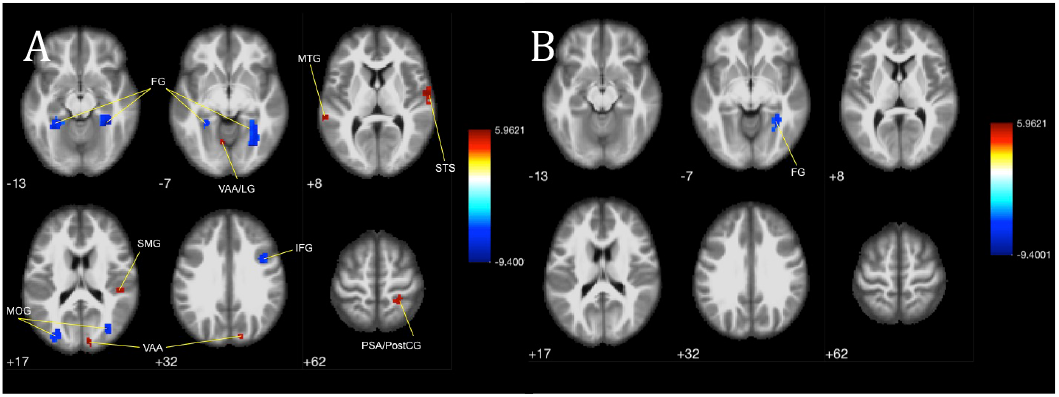
**(A)** Brain activations to typical ethnicities in Singapore in cultural versus ethic context (typical faces in CulCon > typical faces in EthCon), shows activation in multiple regions compared to (**B)** which shows activation only in the FG for non-typical ethnicities in cultural versus ethnic context (non-typical faces in CulCon > non-typical faces in EthCon).

**Table 2.**
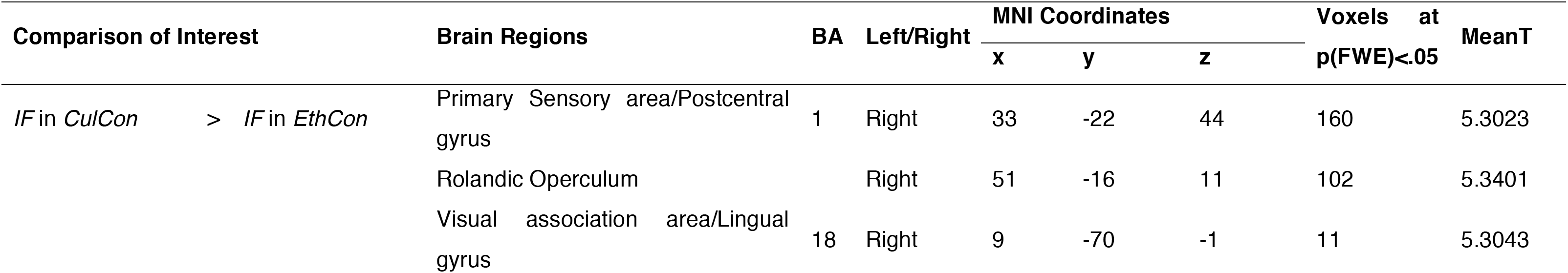
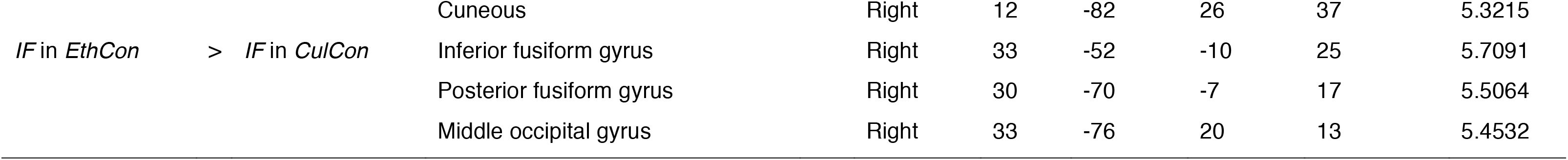
Sites of significant neural activation (p < 0.05. FWE correction) related to the *IF* in *CuiCon* vs. *IF* in *EthCon*

Peaks of all brain activations for these respective contrasts are summarized in Table 1. Refer to Figure 2 for the difference in activation patterns for faces of typical ethnicities and non-typical ethnicities in ethnic and cultural contexts.

### Neural Responses to In-group Faces in Cultural and Ethnic Contexts

To investigate brain responses to in-group Chinese faces in cultural and ethnic contexts, we compared neural responses of in-group faces in the cultural context condition to in-group faces in the ethnic context condition; in-group Chinese faces (IF) in the cultural (CulCon) *versus* the ethnic (EthCon) context conditions (contrasts 2.1). We found significant activated clusters as follows:

– *IF in CulCon > IF in EthCon:* Significant activations of the right VAA comprising of the LG, right PSA comprising of the postCG, and rolandic operculum (RO), a part of the secondary somatosensory area was obtained.
– *IF in EthCon > IF in CulCon:* Higher activations in FG and MOG. All activations were limited to the right hemisphere (Figure 3).

**Figure 3.**
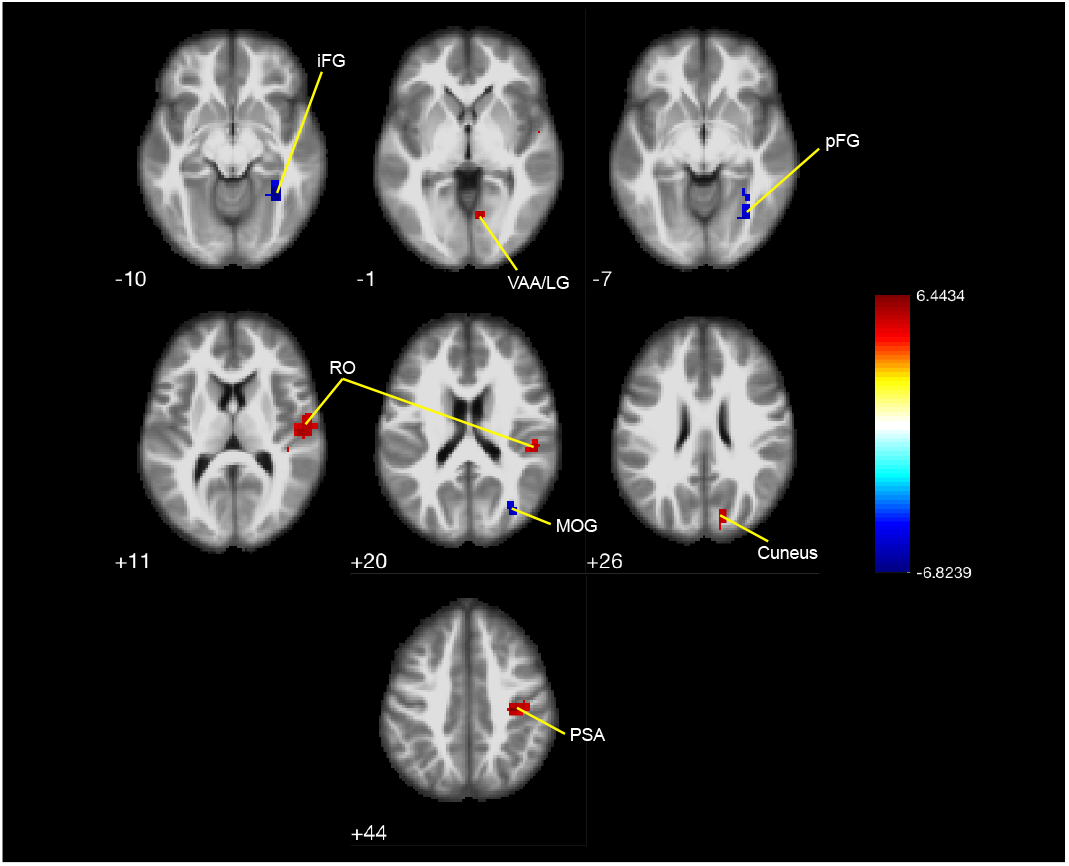
Brain activation of in-group Chinese faces in cultural context versus ethnic context (IF in CulCon > IF in EthCon).

### Neural Responses to Typical Out-group (typical other-race) Faces in Cultural and Ethnical Contexts

Neural responses of culturally primed out-group faces were compared to ethnically primed out-group faces; ***2.2) OF in CulCon vs. EthCon**,* to understand the effect of context on out-group faces. Due to the lack of ample trials, as we have obtained this contrast from a larger task, and with a stringent FWE correction, no significant activation was found at p(FWE)<0.05. However, with a less stringent uncorrected p-value< 0.001 we found significant cerebral brain activation in (Table 3):

− *OF in CulCon > OF in EthCon:* Higher activation in the VAA, right FG, and the left postCG, a part of the somatosensory cortex.

**Table 3.** Sites of significant neural activation (p< 0.001. uncorrected) related to the contrast *OF* in *CulCon* vs. *OF* in *EthCon.*

| Comparison of Interest | | | Brain Regions | BA | Left/Right | MNI Coordinates | | | Voxels at<br>$p(\text{uncorr}) < .0001$ | MeanT |
| --- | --- | --- | --- | --- | --- | --- | --- | --- | --- | --- |
|  |  |  |  |  |  | x | y | z |  |  |
| *OF* in *CulCon* | > | *OF* in *EthCon* | No voxles survied threshold |  |  |  |  |  |  |  |
| *OF* in *EthCon* | > | *OF* in *CulCon* | Fusiform |  | Right | 33 | -46 | -7 | 107 | 3.9091 |
|  |  |  | Lingual |  | Left | -24 | -64 | -7 | 74 | 3.7296 |
|  |  |  | Visual association area | 18 | Right | 15 | -73 | -1 | 23 | 3.5755 |
|  |  |  | Postcentral gyrus |  | Left | -39 | -19 | 26 | 18 | 3.4173 |

### Preliminary contrasts for Difference and Similarities of Neural Responses to Out-group and In-group based on Context

#### Comparing Neural Responses of Out-group and In-group in Cultural Context

If a common shared culture does drive an automatic and spontaneous sense of membership to a multi-ethnic group, then brain responses to out-group faces primed with culture, representative of an enlarged cultural in-group, should be similar to brain responses to in-group faces. Thus, any differences between out-groups and in-groups in the cultural context were assessed; **2.3) *OF vs. IF in CulCon***.

− *OF in CulCon > IF in CulCon:* We found no significant differences in brain responses to in-group and out-group faces in the cultural context at p(FWE) < 0.05 (Figure 4A). Even at a less stringent uncorrected p<0.001, the only difference was a higher activation in the right FG when participants viewed in-group compared to out-group faces (see Table 4).

**Figure 4.**
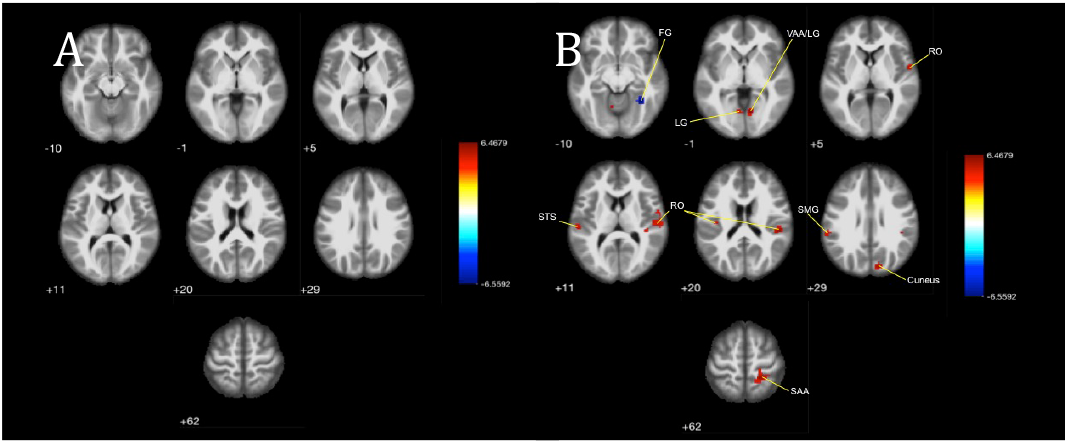
**(A)** No significant difference in brain activation when viewing out-group faces versus in-group faces in the cultural context (OF in CulCon > IF in CulCon) suggesting an enlarged in-group based on cultural identity, which is not present in **(B)** when viewing out-group versus in-group faces in the ethnic context (OF in EthCon > IF in EthCon).

**Table 4.** Sites of significant neural activation related to the contrast *OF* in *CulCon* vs. *IF* in *CulCon.*

| Comparison of Interest | Brain Regions | BA | Left/Right | MNI Coordinates |  |  | Voxels | MeanT |
| --- | --- | --- | --- | --- | --- | --- | --- | --- |
|  |  |  |  | x | y | z |  |  |
| *p*(FWE) < 0.05 |  |  |  |  |  |  |  |  |
| *IF* in *CulCon* > *OF* in *CulCon* | - |  |  |  |  |  |  |  |
| *OF* in *CulCon* > *IF* in *CulCon* | - |  |  |  |  |  |  |  |
| *p*(uncorr) < 0.0001 |  |  |  |  |  |  |  |  |
| *IF* in *CulCon* > *OF* in *CulCon* | Fusiform gyrus |  | Right | 33 | -46 | -7 | 42 | 4.5133 |
| *OF* in *CulCon* > *IF* in *CulCon* | - |  |  |  |  |  |  |  |

#### Comparing Neural Responses of Out-group and In-group in Ethnic Context

The contrast **2.4) *OF vs. IF in EthCon*** was examined to compare how out-group and in-group faces were processed in the ethnic context.

− *OF in EthCon > IF in EthCon:* Higher activation of the right VAA extending to the LG, left LG, right sensory association area (SAA), right cuneus, RO a part of the secondary somatosensory area, left STG, and left SMG were found when processing out-group faces (see Table 5 and figure 4).

**Table 5.** Sites of significant neural activation (p< 0.05. FWE) related to the contrast *OF* in *EthCon* vs. *IF* in *EthCon.*

| Comparison of Interest | | Brain Regions | BA | Left/Right | MNI Coordinates | | | Voxels at<br>$p(\text{FWE}) < .05$ | MeanT |
| --- | --- | --- | --- | --- | --- | --- | --- | --- | --- |
|  |  |  |  |  | x | y | z |  |  |
| *IF* in *EthCon* | > *OF* in *EthCon* | Fusiform area |  | Right | 33 | -52 | -10 | 35 | 5.4098 |
| *OF* in *EthCon* | > *IF* in *EthCon* | Visual association area/Lingual gyrus | 18 | Right | 9 | -70 | -1 | 32 | 5.4935 |
|  |  | Lingual gyrus |  | Left | -9 | -64 | -7 | 27 | 5.5316 |
|  |  | Sensory Association area | 5 | Right | 21 | -43 | 62 | 98 | 5.2934 |
|  |  | Rolandic Operculum |  | Right | 51 | -31 | 20 | 89 | 5.2849 |
|  |  | Cuneus |  | Right | 12 | -82 | 29 | 34 | 5.3759 |
|  |  | Rolandic Operculum |  | Left | -39 | -19 | 23 | 10 | 5.2938 |
|  |  | Rolandic Operculum | 6 | Right | 54 | -4 | 5 | 21 | 5.2983 |
|  |  | Superior Temporal gyrus |  | Left | -57 | -25 | 11 | 18 | 5.2113 |
|  |  | Supramarginal gyrus | 40 | Left | -60 | -34 | 29 | 12 | 5.1392 |

### The Difference and Similarities of Neural Responses to Out-group and In-group based on Context

*OF in CulCon vs. IF in CulCon* differed from *OF in EthCon vs. IF in EthCon.* Similarities in neural responses of these contrasts were also assessed through conjunction analysis. Table 6 displays both similarities and differences in activations for the abovementioned contrasts. The cortical midline regions were significantly more active for the cultural context condition compared to the ethnic context condition. These regions were the left and right lingual gyrus, right precuneus and cuneus, and left precentral gyrus. On the contrary, the left and the right fusiform gyrus were significantly active for both cultural and ethnic context conditions.

**Table 6.** Sites of significant neural activation (p< 0.05. FWE) related to the contrast *IF* in *CuiCon* > *OF* in *CuiCon* vs. *IF EthCon* > *OF* in *EthCon.*

| Comparison of Interest | Brain Regions | BA | Left/Right | MNI Coordinates |  |  | Voxels at<br>q(FWE)<.05 | MeanT |
| --- | --- | --- | --- | --- | --- | --- | --- | --- |
|  |  |  |  | x | y | z |  |  |
| *IF* in *CulCon* ><br>*OF* in *CulCon* ><br>*IF* in *EthCon* ><br>*OF* in *EthCon* | Lingual gyrus |  | Right | 9 | -70 | -1 | 42 | 5.3767 |
|  | Postcentral gyrus/Precuneus | 7 | Right | 18 | -45 | 65 | 73 | 4.9517 |
|  | Cuneus |  | Right | 12 | -85 | 29 | 34 | 5.0007 |
|  | Lingual gyrus |  | Left | -9 | -73 | -4 | 10 | 4.9263 |
|  | Precentral gyrus |  | Left | -36 | -28 | 62 | 29 | 4.8493 |
| *IF* in *EthCon* ><br>*OF* in *EthCon* ><br>*IF* in *CulCon* ><br>*OF* in *CulCon* | - |  |  |  |  |  |  |  |
| Conjunction analysis | Fusiform |  | Right | 33 | -52 | -10 | 132 | 5.8720 |
|  | Fusiform |  | Left | -30 | -43 | -13 | 36 | 5.1782 |

## Discussion

This study postulated that by identifying with a common shared culture among various racial ethnicities in multicultural societies, the neural responses to other- and same-race faces would be similar, indicating an automatic and spontaneous sense of membership to multi-ethnic groups (that we called *enlarged cultural in-group effect*). Specifically, using a modified priming paradigm, we investigated the neural response to out-group faces of typical ethnicities present in Singapore (Indian faces), non-typical ethnicities (Arab and Caucasian faces) and in-group faces (Chinese) in the ethnic and in the (shared) cultural context conditions.

### More effort in processing out-group faces in ethnic context contrasted to cultural context

When typical out-group faces (typical other-race face; Indian) were viewed in the ethnic as compared to the cultural context conditions, a high engagement of visual processing and social cognitive regions was found. Specifically, the ethnic context increased the cerebral activation in the VAA, implicated in enhanced visual processing (Rosen et al., 2018); in the lingual gyrus, associated with outward-directed social and contextual focus (Colton et al., 2013); and in the left postCG, part of the somatosensory cortex (Chan & Baker, 2015; Green et al., 2018), involved in mentalizing networks, social emotions and social perception (Keysers et al., 2010; Sugranyes et al., 2011; Bernhardt & Singer, 2012; Schaefer et al., 2013; Valchev et al., 2016). Activation in the FG was also present. The FG is implicated in top-down and bottom-up perception of visual input. Bottom-up perception includes higher FG activation to racial in-group faces than out-group faces (Guassi Moreira et al., 2016; Molenberghs, 2013; Shkurko, 2012; Van Bavel et al., 2008). The top-down modulation of FG in social perception involves indexing socially significant perceptual input according to preexisting schemas of the social environment (Shkurko, 2012). Thus, FG is considered of the first few regions to respond to social faces and may represent the top-down process involved in explicit social categorization in response to the social context (Shkurko, 2010). Therefore, activations of higher visual processing and social cognitive regions for out-groups in the ethnic context may indicate more effort in processing the out-group faces in the ethnic context.

Brain regions characteristic of out-group activation such as the amygdala (Derntl et al., 2009; Freeman et al., 2010; Molenberghs, 2013; Shkurko, 2010; Van Bavel et al., 2008), ACC (Cunningham et al., 2004; Kaplan et al., 2007; Knutson et al., 2007), or insula (Falk et al., 2012; Kaplan et al., 2007; Phelps et al., 2000) were not found in the present study. Amygdala is implicated in social categorization and implicit racial bias (Cunningham et al., 2004; Shkurko, 2010). Cunningham et al. (2004) reported amygdala activation when faces were presented before conscious processing (30ms) could occur. Such activation was absent when faces were presented for a longer duration (525ms), allowing for conscious processing where implicit racial bias would be regulated. Therefore, a possible reason for the lack of amygdala activation could be because the face stimuli were presented for 4 seconds, allowing for any regulation of racial bias. If this stands true for the current study, then ACC activation should have been obtained, to consciously regulate out-group bias, similar to that of Cunningham et al.’s findings (2004). However, no activation in the ACC was found here. Zuo and Han (2013) reported that neural activity in the pain matrix, including the anterior cingulate cortex and anterior insula, did not differ for Asian and Caucasian models when Chinese adults, brought up in Western countries, were made to see video clips of either Asian or Caucasian models receiving painful stimulations. Derntl et al. (2009) reported a decrease in amygdala activation in Asians who stayed longer in Europe during an explicit emotion recognition task involving Caucasian actors. Therefore, it is possible that exposure to multi-ethnic societies, including out-groups, generally reduced racial bias.

### Typical ethnicity and same-race faces when primed by ethnic context contrasted to cultural context

Brain regions involved in social cognition, self-perception networks, and theory of mind networks were less activated in the ethnical context, than the cultural context. When out- and in-group faces of typical ethnicities (Chinese, Indian) were viewed in the ethnic context, contrasted to cultural contexts, higher activation of the FG was obtained, which is one of the core brain regions for face recognition (Tovée, 1995; Zhen et al., 2013). FG is commonly associated with in-group face processing (Guassi Moreira et al., 2016; Molenberghs, 2013; Shkurko, 2012; Van Bavel et al., 2008). However, there are studies that have reported, similarly, FG activations during out-group face processing (Ronquillo, 2010; Shkurko, 2010; Iidaka, 2014). Although the MOG is involved in initial face processing as well (Pizzagalli et al., 2000), it also forms connections to higher social cognition areas such as the STS (Duvernoy & Bourgouin, 1999; Jiang et al., 2018) and specialized cortices involved in emotional and value processing (Olson et al., 2013). Certain connections to higher-level social cognitive areas may be due to the mixture of in-group Chinese and out-group Indian faces in typical ethnicities. A subset of similar regions was activated for in-group faces in the ethnic context.

However, same-race faces (in-group) in the cultural context, compared to ethnic context, elicited higher activation in mainly the MNS regions which are brain regions involved in social perception, and includes the VAA comprising of the lingual gyrus (Adolphs, 1999; Colton et al., 2013; Freeman et al., 2010; Scheepers et al., 2013), and the TPJ networks. The MNS system is famous for its activation while individuals observe the behavior of other people (Rizzolatti & Craighero, 2004), and implicated in neurocognitive functions such as social cognition, empathy, and theory of mind (Rizzolatti & Craighero, 2004; Sugranyes et al., 2011). TPJ is associated with the theory of mind and mentalizing (Freeman et al., 2010; Shkurko, 2012). Theory of mind refers to the process of explaining or predicting other people’s behavior by way of reading, reasoning, and representing their independent mental states (Freeman et al., 2010). These regions are involved in self-other distinctions and making inferences about people (Greven & Ramsey, 2017; Molenberghs, 2013). Consistent with the in-group findings, typical ethnicity faces in the cultural context, compared to ethnic context, elicited higher activations in regions associated with social perception and cognition (STS; MTG, VAA to the lingual gyrus, along with the MNS and the PSA).

Interestingly, typical ethnicities and non-typical ethnicities were processed differently in both ethnic and cultural contexts. We did not observe any context effect when atypical other-race faces were viewed in the ethnic context, compared to the cultural context. Participants showed more sensitivity to contextual primes for typical ethnicities (Chinese and Indian faces) as opposed to non-typical ethnicities (Arab and Caucasian faces). One explanation is that our experimental design primarily tested the effect of a common shared culture between different ethnic groups, that also share common values of the belonging society. As a check on typical faces of different ethnicity, we added faces of people of diverse and non-typical ethnicity. In this sense, the results support the idea that manipulation of the context loses its meaning in the category of atypical ethnicity as there is no mutual cultural sharing.

Moreover, the present study showed that high social cognitive regions were recruited while viewing faces in the cultural context compared to more automatic and implicit processing of faces when viewed in their respective ethnic contexts. We found that viewing of in-group faces in the ethnic context showed typical in-group face brain activation compared to viewing them in the cultural context (i.e., higher activations in the FG (Iidaka, 2014; Molenberghs, 2013; Van Bavel et al., 2008) and MOG (Jiang et al., 2018; Olson et al., 2013).

Overall, results indicate that in the cultural context, individuals engage higher social cognitive resources in order to make individuating judgments of people rather than superficial judgments based on group categorization (Freeman et al., 2010).

### Enlarged In-group Present in Cultural Context

When neural responses to typical other- (out-group) and same-race (in-group) faces were compared in the cultural context, no significant difference was found. Even using the less stringent p-value (uncorrected p<0.0001), the only difference was activation in the right FG for in-group faces. In contrast, the difference between the out-group and in-group faces in ethnic contexts was much more significant. Greater effort in social, cognitive, and affective processing was found for out-group faces in the ethnic context, as discussed earlier. As expected, the lack of difference in neural responses to out-group (typical other-race) and in-group faces indicates that out-groups are processed similarly to in-groups. This phenomenon is characteristic of an enlarged cultural in-group. These findings are similar to the pattern of findings by Zuo, and Han (2013), who found no difference in neural activity in the pain matrix for in-group (Asians) and out-groups (Caucasians) amongst Chinese adults brought up in Western countries.

Specifically, the contextual manipulation of culture in this study may only be relevant to prime a common shared culture that is bound to exist only amongst well-integrated typical ethnicities, which may have rendered the contextual manipulation insensitive for non-typical ethnicities as seen in this study.

The analysis conducted to assess the difference in how neural responses differ for in-groups and out-groups based on context showed higher activation in the midline cortical regions for the cultural context compared to the ethnic context. This further suggests that the out-groups and in-groups may be processed similarly in the cultural context owing to individuating judgments of people rather than superficial judgments based on group categorization (Freeman et al., 2010), as suggested by high midline cortical activation in the cultural context.

Findings from this study suggest that identifying with a common shared culture, more than ethnicity, can drive a spontaneous sense of membership to a multi-ethnic group when the society explicitly and positively supports cross-ethnic interactions in every daily life context. These findings demonstrate that living in proximity and with an inclusive attitude with citizens from other ethnicities can expand the sense of membership of different individuals to that of a member of the same cultural group. The results shed light on a possible emergent phenomenon of a new way to categorize oneself and others, in terms of membership, which stands between the classical ethnic in- and out-group categorization and where culture and not ethnicity shapes and drives spontaneous social judgment of others (Figure 5).

**Figure 5.**
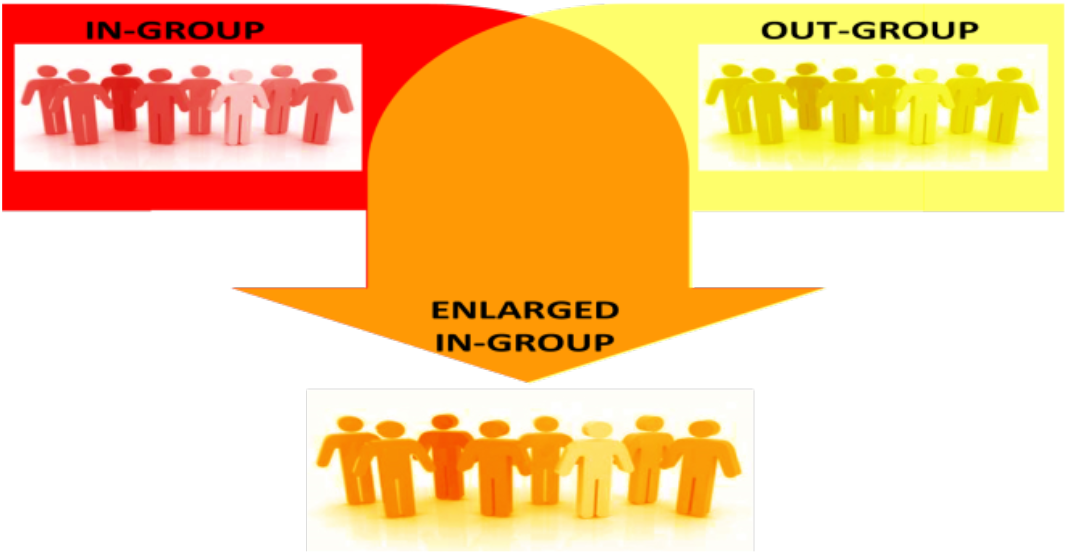
Graphical representation of a likely scenario of in- and out-group membership dynamics in Singapore. Red and yellow colors are namely ethnic-based in-groups and out-groups; orange color is the new phenomenon of the culture-based enlarged in-group which integrates features from both of the above mentioned groups.

However, further research can also be extended to investigating brain reactivity and responsiveness to different ethnic baby faces in a multicultural society and the effect of the environmental context. Such insight will give a substantial theoretical contribution to building an integrated and general model of the social brain.

## Limitations

Firstly, only the majority ethnicity of Singapore (Chinese) participants participated in the study. However, majority and minority groups may perceive and experience intergroup relations differently following their experiences (Dovidio et al., 2008). In order to get a holistic understanding, neural responses to enlarged cultural in-groups should be investigated amongst minority groups as well.

Another limitation is the sample of the age group. All participants were young adults (students or recently graduated). Age differences were reported in studies involving in-group and out-group (Guassi Moreira et al., 2016) and social cognition areas (Colten et al., 2013). Therefore, the effect of enlarged cultural in-group is necessary to be investigated across generations to generalize the findings wholly.

Posing another limitation is that the face stimuli used in this study are only of the female gender. Research from intergroup relations has shown that gender influences in-group racial bias (e.g., Navarrete et al., 2010; Rudman & Goodwin, 2004). Face stimuli should be both male and female in order to analyze any gender effects.

Lastly, all the faces were unfamiliar to participants. Familiar and new faces have differing neural activation. Intergroup friendships are common in multicultural societies. Therefore, it is necessary to understand the neural representation of familiar faces in ethnic in-group, culturally enlarged in-group, and out-group faces.

## Conclusion

Although findings from this study come with certain limitations, it is the first step towards integrating and constructing a model of the multicultural social brain. Building and identifying with a common shared culture in multicultural societies enhance the individuating processing of ethnic out-groups and leads to the formation of an enlarged cultural multi-ethnic in-group. In a well-integrated multicultural society, members do not innately respond to other ethnicities with typical out-group bias responses, but dispense more social and cognitive resources into perceiving them more as individuals than superficially as an out-group. These pioneering findings of the existence of an enlarged cultural in-group open the opportunity for future research to develop on the understanding of multicultural societies and its complex interaction with contextual environment towards dynamic perceptions of in-groups and out-groups through a multi-level approach.

## Acknowledgements

All participants in this study are gratefully acknowledged. The authors thank Lim Mengyu for assistance. This research was supported by NAP SUG 2015 (GE) and Singapore Ministry of Education ACR Tier 1 (GE; RG149/16 and RT10/19).

